# The fitness landscape of a Form II rubisco in a photosynthetic bacterium guides engineering of oxygen tolerance

**DOI:** 10.64898/2025.12.07.690893

**Authors:** Ute A. Hoffmann, Axel Knave, Anna Z. Schuppe, Sebastià Capó-Bauçà, Karen Schriever, Emil Sporre, Rui Miao, Jeroni Galmés, Per-Olof Syrén, Elton P. Hudson

## Abstract

Rubisco is an important but challenging protein engineering target. Fast and selective rubiscos could enhance photosynthesis in plants and accelerate biobased production processes. To facilitate engineering of rubisco, we applied an in vivo screen that couples rubisco activity to growth rate of the photoautotrophic cyanobacterium Synechocystis sp. PCC 6803. We screened a barcoded mutagenesis library of the form II rubisco from Gallionella sp. containing 15,000 single-site and multi-site variants. Exchanges in loop 6 near the active site, at the dimer interface, and in potential gas tunnels improved rubisco fitness. The dataset also informed protein engineering, using recombination and a trained transformer model. In vitro characterisation of two high-fitness variants showed reduced catalytic efficiency for oxygenation (k*_cat_*/K_o_) in both and an increased carboxylation turnover (*k_cat_*^C^) in one. This large labeled fitness dataset, containing examples of epistasis, can be useful for benchmarking computational models of rubisco.

**Teaser:** Evolving a foreign rubisco to photosynthesis leads to reduced oxygen sensitivity.

## Introduction

Rubisco is an important engineering target for improving photosynthetic efficiency in plants (1,2). Enhancing the specific activity of rubisco could reduce nitrogen demand, while enhancing rubisco selectivity for CO2 over O2 could reduce the accumulation of 2- phosphoglycolate (2-PG), a byproduct that inhibits carbon fixation and must be salvaged by photorespiration (3,4). Additionally, rubisco may limit carbon conversion in certain microbial bioproduction platforms, such as cyanobacteria and lithoautotrophic bacteria (5). However, protein engineering of rubisco is challenging. Expression of plant rubiscos in bacteria requires multiple maturation factors and chaperones (6). The observed correlation between carboxylation and oxygenation catalytic efficiencies in rubiscos (7,8) may be indicative of a mechanistic coupling (9), though it was proposed that plant rubiscos are evolving to increase carboxylation efficiency and selectivity (10,11). Bacterial rubiscos have been pursued as replacements for the plant enzyme in crops, as these are expressed at higher titers, and generally display higher carboxylation rates, though their lower selectivity necessitates a high CO2 cultivation (12–16). There has been some success in enhancing the kinetic parameters of bacterial form I rubiscos, such as increasing carbon fixation rate or reducing oxygen sensitivity, though rarely were these two achieved simultaneously (17–20). Form II rubiscos represent an alternative rubisco evolutionary lineage, primarily adapted to low O2 environments, and include the highest carboxylation rates reported (21–23). The simple format of form II rubiscos, which lack small subunits (24), may increase the available space for mutation since conservation of contacts to other proteins are not required (25–27). Form II rubiscos have also been subjected to extensive mutagenesis in order to map the determinants of speed and selectivity (28–31). This literature suggests that some kinetic parameters can be modulated individually. For example, the low CO2 affinity of a form II rubisco could be improved by single point mutations without an apparent tradeoff in carboxylation rate (29,32). Recently, it was reported that the binding affinity of form II rubisco from *Gallionella* for O2 could be reduced with little effect on the carboxylation efficiency, resulting in rubisco mutants that had higher catalytic efficiency in air than the wild type, and thus imparted better growth to *E. coli* in aerobic conditions (22). Form II rubiscos with high carboxylation efficiencies and reduced sensitivity to oxygen could be starting points for eventual transfer to plant or other bioproduction systems.

*In vivo* screening platforms for rubisco activity expand the number of variants that can be tested simultaneously. Such platforms can also be adapted in order to assay specific kinetic parameters. For example, changes in rubisco specificity factor (*S*C/O) can be estimated by comparing enrichment of mutant clones during oxic or anoxic growth, or CO2 affinity by growth at low CO2. To date, rubisco-dependent *E. coli* screens have relied on error-prone PCR (Ep-PCR) to generate enzyme diversity, and coupled these libraries to directed evolution schemes to enrich high-fitness variants (33). Ep-PCR generates multi-site mutants, which allows study of epistatic relationships in a protein (25,34–36). However, Ep-PCR results in a large number of inactive enzymes. Furthermore, much of the information on enzyme fitness is lost, as less fit variants are not retained or characterized further. To overcome the first limitation, input libraries can be designed using prior knowledge, such as phylogenetic relationships (37). Using prior knowledge has a higher likelihood of identifying a more fit variant than random input (37,38). To overcome the second limitation, variants can be barcoded and tracked using deep sequencing, such as in deep mutational scanning (39). Labeled fitness data, often consisting of thousands of data points, can be used to train machine-learning models to eventually guide precise engineering for enzyme catalytic parameters (40).

Recently, we established a rubisco screening strain of the cyanobacterium *Synechocystis* sp. PCC 6803 and demonstrated that the platform could be used to screen activity of the form II rubisco from *Gallionella* sp., CbbM (41). A closer evolutionary relationship between plants and cyanobacteria may allow us to select for rubisco properties that are relevant for photoautotrophy, such as improved resistance to *in situ* generated oxygen, or resistance to inhibition from prevalent sugar phosphates. Furthermore, faster rubiscos could be leveraged for promoting productivity of cyanobacteria in bioprocesses (42,43). We also established a combination of the phylogenetically-guided method EVmutation (44,45) and *in silico* evolution (46) to design higher-order rubisco variants to be tested in the cyanobacteria screening strain (41). In this work, we extended this approach and used both EVmutation and DeepSequence (47) to predict fitness values of variants identified by *in silico* evolution. We chose the models EVmutation and DeepSequence due to their comparably high performance in predicting the effect of amino acid exchanges in benchmark datasets, as well as code accessibility (48,49). We identified seven amino acid positions and specific exchanges at these positions, which had, according to both algorithms, a high likelihood to increase Rubisco fitness. This combinatorial library was combined with a deep mutational scanning. The complete library, consisting of saturation/DMS and combinatorial parts, was introduced into the screening strain, and we performed growth selection experiments under different gas feed and light conditions. This resulted in an extended fitness landscape of *Gallionella* Rubisco and was used to create second-generation Rubisco variants. These variants were created both through a naive approach and by using the machine learning algorithm ProteinNPT.

## Results

Recently we reported an *in vivo* screening platform for heterologous rubisco in the photoautotrophic cyanobacterium *Synechocystis* sp. PCC 6803, which we applied to screen a small barcoded mutagenesis library of the rubisco CbbM from *Gallionella* sp. (41). In this strain, the native form I Rubisco is repressed by 99% using inducible CRISPR interference (CRISPRi) and cell growth rate becomes dependent on CbbM activity (Fig. 1A). As CbbM is not encapsulated in the *Synechocystis* carboxysomes, growth rate is also influenced by the CO2 level and the CO2:O2 ratio in the culture gas feed. We reasoned that this screening platform would make it possible to rank mutant rubiscos based on kinetic parameters such as *k*cat^c^, KC, and *S*C/O. In the present work, we extended screening to a large, barcoded library of CbbM variants (Fig. 1A). The starting point was variant CbbM_base, a 2-point mutant of CbbM (S162P, G422R) that we previously found to impart a similar growth to *Synechocystis* as CbbM, but is more thermostable (CbbM_base Tm 50 C, CbbM Tm 44 C) (41). This variant was chosen as base for the library since thermostable enzymes on average tend to tolerate mutations to a higher degree (50).

**Figure 1:**
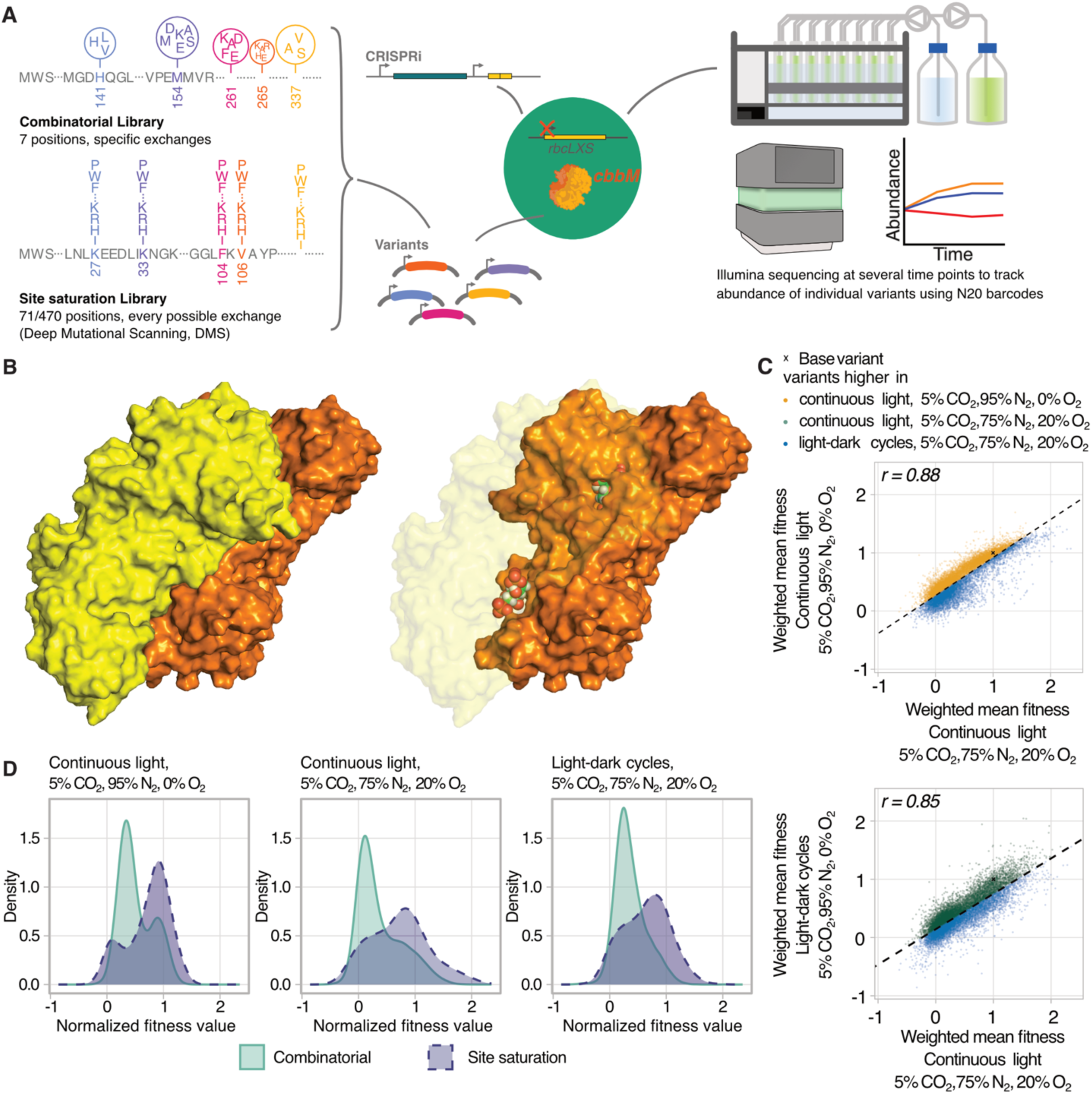
Workflow and fitness distributions of the CbbM library. (**A**). A combinatorial library with selected exchanges at seven positions and a saturational library with single amino acid exchanges were combined and introduced into a *Synechocystis* mutant strain in which the endogenous rubisco can be repressed by CRISPR inhibition (CRISPRi). The pooled *Synechcoystis* was subjected to growth competition in three conditions. White light intensity was 300 µE in all cases. The abundance of individual mutant variants was tracked by Illumina sequencing of assigned 20-nucleotide barcodes. (**B**) Predicted structure of the CbbM WT dimeric enzyme with the CAP (an analog of RuBP) at the active site shown as spheres. (**C**) Comparison of normalized weighted mean fitness values for the complete library at different growth conditions. The dashed line gives the line defined by the linear regression of both conditions. Pearson’s r for the correlation of the respective conditions is 0.88 and 0.85, respectively. (**D**) Density plots of fitness distributions of combinatorial and saturation library parts at different growth conditions. Fitness values were normalized so that the CbbM_base variant had a fitness score of 1.0 and the median of fitness values for inactive K200 variants had a fitness score of 0.0.

### A phylogenetically-guided approach to designing an input library for high-throughput screening

We designed a CbbM_base library comprising both saturation and combinatorial mutagenesis of selected residues. Positions for saturation mutagenesis (totaling 71 positions) were chosen using phylogenetically-guided methods DeepSequence (47) and EVmutation (44,45), which use MSAs to identify residues of high conservation or that may be co-evolving. We picked residue positions for which the EVmutation epistatic score was above 1.0, indicating that exchange at this position could have a positive effect, or which were among the top 100 highest fitness predictions of DeepSequence and also had an EVmutation epistatic score above -1.0 (Supp. Table 1). Additionally, we included 11 lysine residues for saturation mutagenesis, based on the hypothesis that acetylation of these could play a role in enzyme regulation (51). Finally, we also included N389 and N401, positions where mutation was previously shown to have a positive effect on expressibility or catalytic efficiency in rubisco homologues (52,53). Saturation mutagenesis of these 71 positions (15% of CbbM sequence) resulted in 1,349 variants (Supp. Table 2; Supp. Data File 1).

**Table 1:**
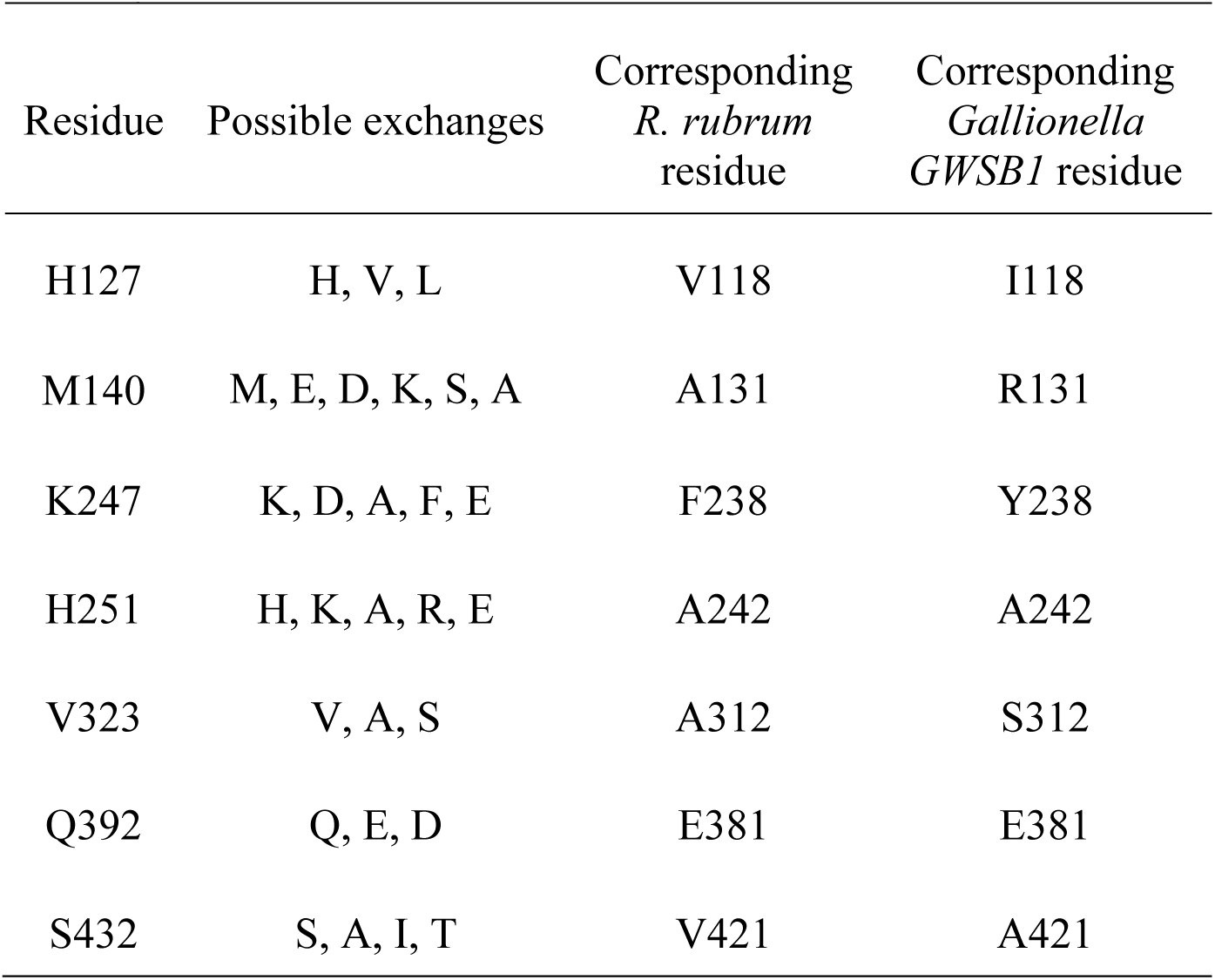
Amino acid positions and possible replacement residues chosen for the CbbM combinatori<u>al library.</u>.

| Residue | Possible exchanges | Corresponding<br>*R. rubrum*<br>residue | Corresponding<br>*Gallionella*<br>*GWSB1* residue |
| --- | --- | --- | --- |
| H127 | H, V, L | V118 | I118 |
| M140 | M, E, D, K, S, A | A131 | R131 |
| K247 | K, D, A, F, E | F238 | Y238 |
| H251 | H, K, A, R, E | A242 | A242 |
| V323 | V, A, S | A312 | S312 |
| Q392 | Q, E, D | E381 | E381 |
| S432 | S, A, I, T | V421 | A421 |

Positions for combinatorial mutagenesis were selected by coupling the output of EVmutation and DeepSequence to iterative rounds of *in silico* optimization, as described previously (46). This analysis identifies subsets of residues in CbbM and samples multiple amino acid exchanges at these residues to find combinations that are potentially fitness enhancing (41). Based on this, we selected seven residues to be mutagenized with a reduced set of amino acid exchanges (Table 1). As expected from a phylogeny-based method, many of the suggested exchanges corresponded to amino acids present in rubisco homologues (Supp. Fig. 1). The reduced set of exchanges were combined in every possible manner, yielding 16,200 possible variants, of which 16,178 contained more than one amino acid exchange (Supp. Table 2). To enable tracking of different protein variants by Illumina sequencing, each gene variant was labeled with a N20 barcode. The CbbM saturation and combinatorial libraries were synthesized by commercial supplier (GeneScript). After transformation of the CbbM library into *E. coli,* barcodes were mapped to variant sequences using PacBio long-read sequencing. The *E. coli* plasmid pool contained >95% of the designed library variants. The plasmid pool was transformed into the *Synechocystis* screening strain. Deep sequencing of the *Synechocystis* pool showed an average of 2-3 barcodes per variant.

### Single-site exchange library contains many rubisco variants with higher fitness

The resultant *Synechocystis* library was pooled and cultivated under three growth conditions in a multicultivator operating in turbidostat mode: continuous light with an oxygen-free gas feed of 5% CO2, 95% N2, 0% O2, continuous light and an oxygen-containing gas feed of 5% CO2, 70% N2, 20% O2, and light-dark cycles in the oxygen-containing gas feed. The average growth rate was dependent on the oxygen content (0.041 h^-1^ in 0% O2 condition and 0.028 h^-1^ in 20% O2 condition), and was significantly slower than wild type *Synechocystis* (0.08 h^-1^ in similar conditions (41,54)). We note that due to *in situ* generated O2 during photosynthesis, a 0% O2 growth condition is not anoxic. Samples were periodically withdrawn from the bioreactors over 12 generations, and barcodes in the population quantified using Illumina sequencing. For each CbbM variant, we calculated a fitness score based on the clone specific growth rate. Fitness scores were normalized for each condition so that fitness of the CbbM_base variant corresponded to 1, and the median fitness of all non-active K200 variants (30) corresponded to 0 (Supp. Fig. 2). Calculated fitness values showed a bimodal distribution (Fig. 1D), implying a selection based on CbbM properties. The fraction of variants that had higher (p<0.05 Wilcoxon Rank test) fitness than CbbM_base was 8% and 0.8% for the saturation and combinatorial libraries, respectively (Supp. Table 3). Statistical significance depends on both effect size and the number of unique barcodes connected to a variant.

**Figure 2.**
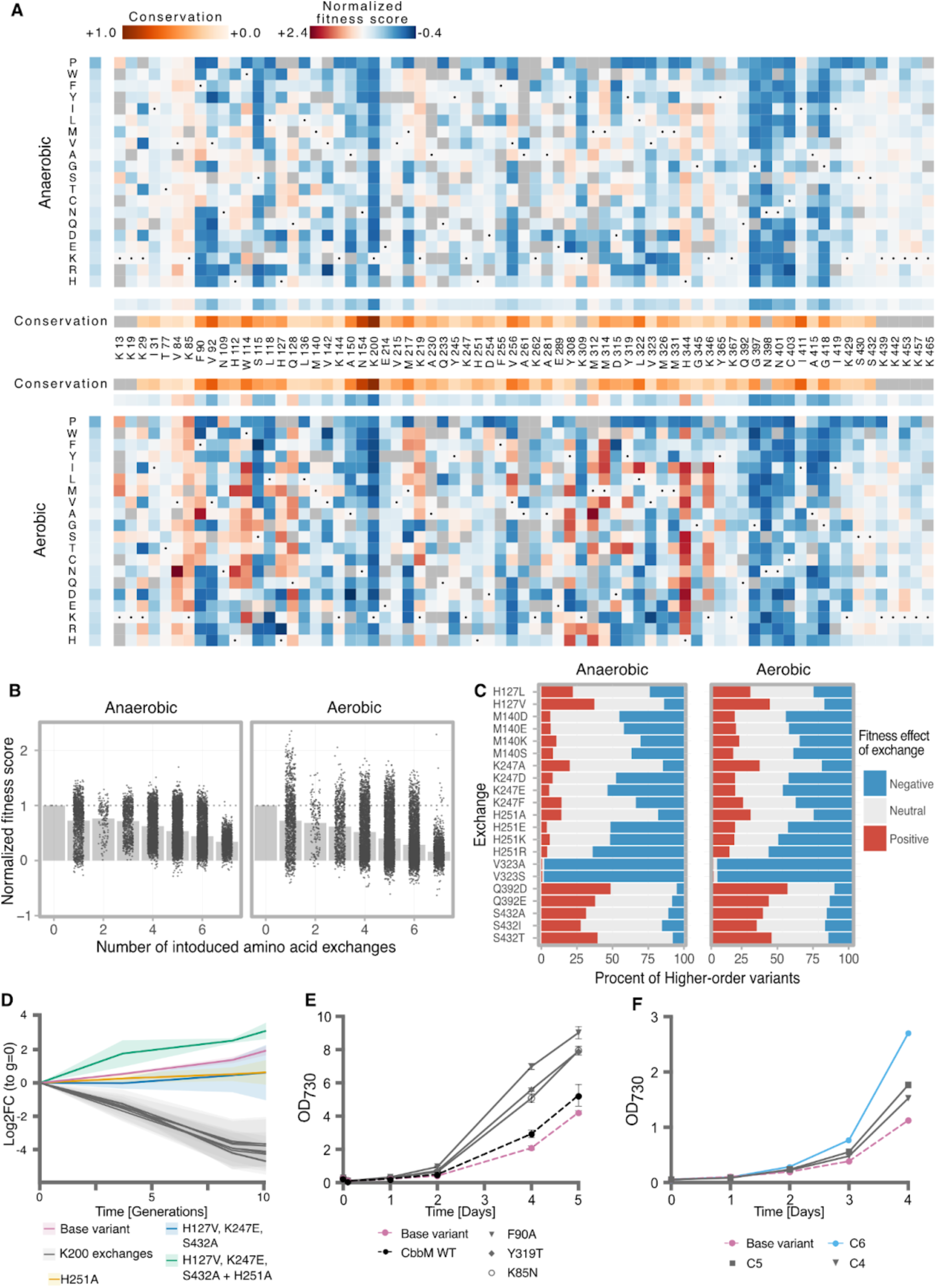
(**A**) Heatmap of variant fitness of the saturation part of the library in two growth conditions. Values are normalized so that neutral mutations have value = 1 and inactive K200 variants have value 0. Gas conditions were 5 % CO2, 95 % N2, 0 % O2, and 5 % CO2, 75 % N2, 20 % O2. Wild-type residues are indicated with a black dot. Gray squares indicate that there is no data available, either because the exchange was not covered by the MSA or because the variant dropped out of the library during construction. (**B**) Normalized fitness scores of the combinatorial library variants plotted against the number of introduced mutations, in nitrogen cultivation and in air condition. Points are fitness scores for variants and bars are averages. (**C**) Global overview of fitness effects of specific amino acid exchanges in either saturation or combinatorial library in condition 5 % CO2, 75 % N2, 20 % O2 . Data for all growth conditions in Supp. Table 4 (**D**) Example of epistasis effect observed for H251A. (**E**) Batch cultivations of selected *Synechocystis* clones from saturation library. Starting OD730 was 0.1 and conditions were 5 % CO2, 75 % N2, 20 % O2 and 50 µE. (**F**) Batch cultivations of selected *Synechocystis* clones from combinatorial library. Starting OD730 was 0.05 and conditions were 5 % CO2, 75 % N2, 20 % O2 and 50 µE.

Fitness scores were negatively correlated with residue conservation score (average Pearson’s r = -0.52; Supp. Fig. 3), though there were non-conserved residues for which all replacements had a detrimental effect (G397, N398, and N401), and a few highly conserved residues were surprisingly tolerant to exchanges (I411, W114). A recent deep mutational scan of *R. rubrum* rubisco reported that solvent exposed residues were more tolerant to mutation than buried residues (29). The fitness of CbbM variants were moderately correlated (average Pearson’s r = 0.47; Supp. Fig. 3) with solvent accessibility.

**Figure 3:**
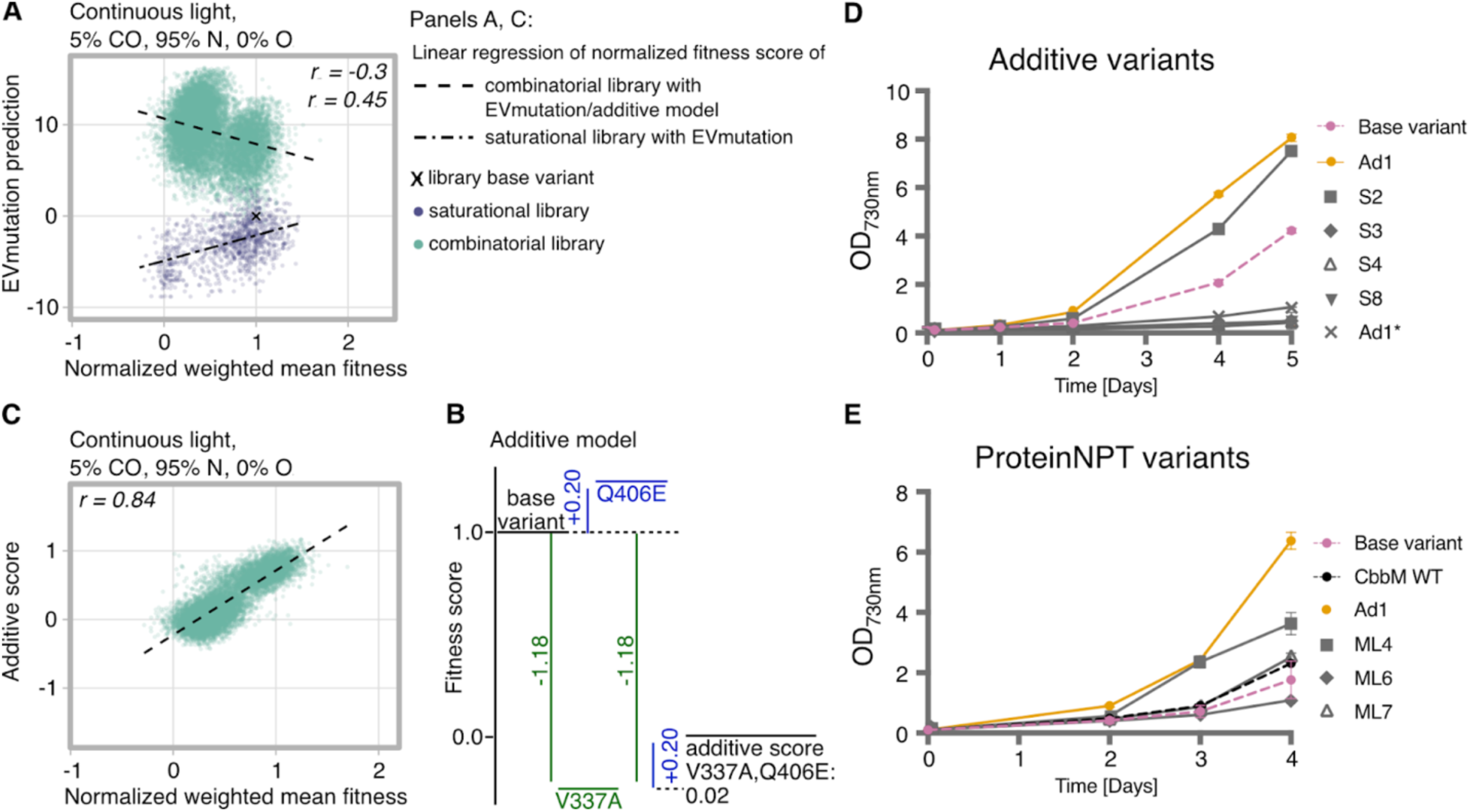
Using labeled fitness data to design CbbM variants. (**A**) Comparison of EVmutation predictions and normalised fitness scores for single-site variants and combinatorial variants. (**B**) Example of additive model for predicting fitness of multi-site variants. (**C**) Comparison of additive model predictions to the normalised fitness scores of combinatorial variants. (**D**) Batch growth of selected second-generation CbbM Synechocystis clones created by the additive method at 5 % CO2, 75 % N2, 20% O2 (**E**) Growth of selected second-generation CbbM Synechocystis clones created by the ProteinNPT method at 5 % CO2, 75 % N2, 20% O2.

Fitness at 21% and 0% O2 growth conditions strongly correlated (Fig. 1C, Pearson’s r > 0.8), though generally fitness increases were higher in the 21% O2 growth condition. Proteomic analysis of the pooled library revealed that *Synechocystis* expressing CbbM has a proteome consistent with carbon limitation, even when grown with a low O2, high CO2 gas feed (Supp. Fig. 4 and Supp. Table 4), which may be a result of photorespiration occurring from *in situ* generated O2. Therefore, it is likely that screening in a O2-free gas feed also imposes a selection pressure on rubiscos with reduced oxygenation activity. Fitness scores of CbbM variants were also highly correlated between constant light and light-dark growth (Fig. 1C). In plants, multiple sugar phosphates have been shown to inhibit Rubisco activity (55). However, Rubisco inactivation in the dark in *Synechocystis* was recently shown to be regulated by RuBP availability and not enzyme inactivation by metabolites (56).

**Figure 4:**
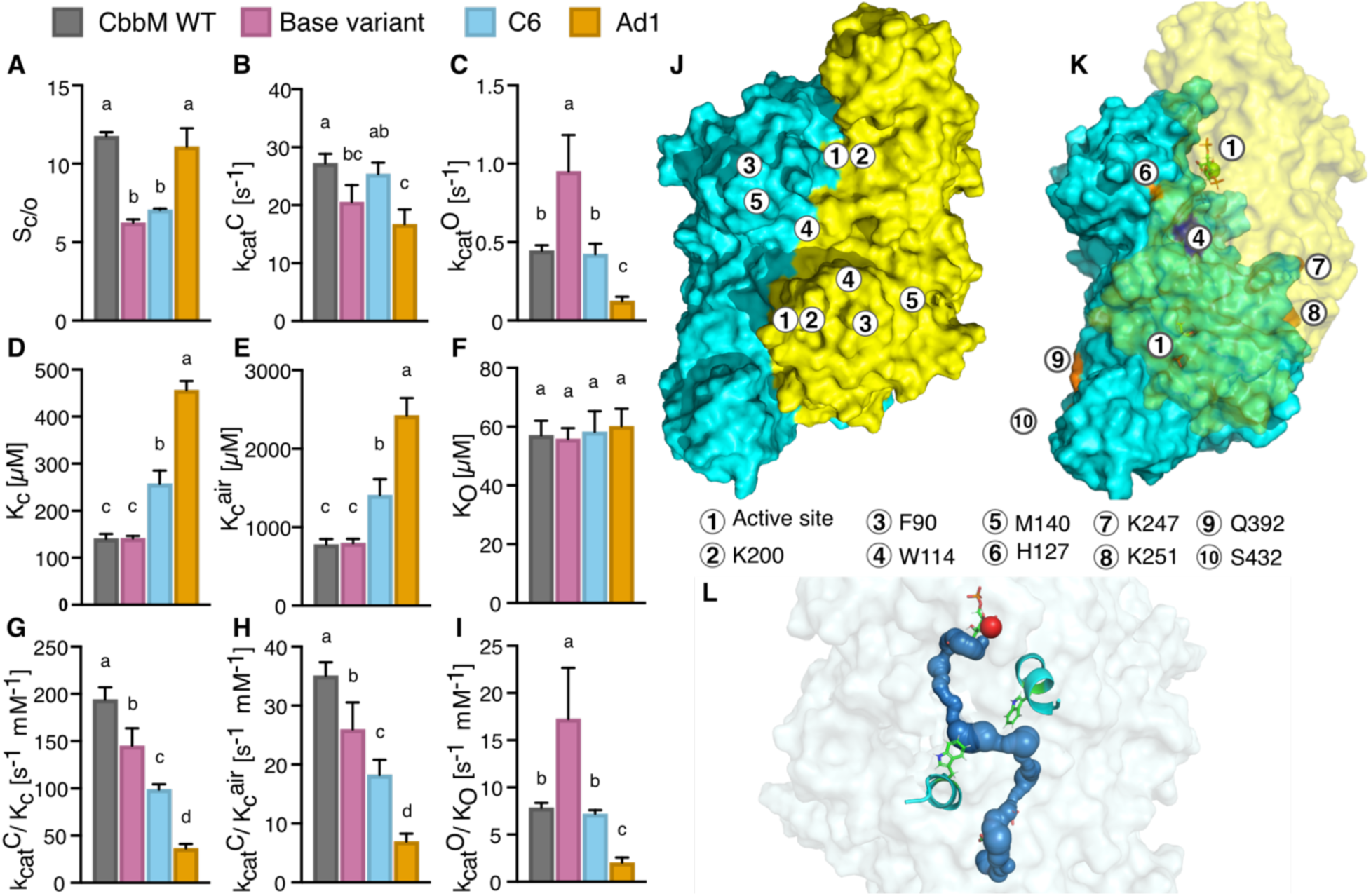
In vitro determined catalytic parameters of select CbbM variants. CbbM_base: S162P, G422R, C6: H127V, M140A, K247A, H251A, Q392E, S432T + S162P, G422R, Ad1: H127V, K247A, H251A, Q392E, S432T, W114I + S162P, G422R. (**A**) Selectivity values of selected CbbM variants. **(B**) Carboxylation rate constant (*k*cat^C^). (**C**) Oxygenation rate constant (*k*cat^O^). (**D**) Binding constant of CO2 in absence of oxygen, (**E**) of CO2 in air, (**F**) binding constant of O2. (**G**) Catalytic efficiency of carboxylation reaction in absence of air, (**H**) of carboxylation reaction in air, (**I**) catalytic efficiency of oxygenation reaction. All catalytic parameters were determined using a ^14^CO2 fixation assay with four replicates. Statistical significance was assessed using a one-way ANOVA followed by Tukey’s multiple comparisons test (**J**) Visualization of important residues and active site in a CbbM model (AlphaFold). The dimers are shown in blue and yellow (**K**) Visualization of CbbM model displaying the location of residues mutated in variant Ad1. (**L**) Residue W114 (green) in CbbM constricts a tunnel (blue) in the dimer interface, connecting the two active sites.

Replacements at a few positions had, on average, a positive effect on the normalized fitness value. Notable were residues H344 and K346, located within the C-terminal hinge of loop 6 which is proposed to be involved in CO2/O2 discrimination and catalysis (31,57–59). Most exchanges at H344 had a positive effect on fitness, except H344G and H344P, which had a negative effect (Fig. 2A). It is tempting to assume that these latter exchanges destabilize the loop by affecting backbone flexibility. Interestingly, exchanges H344W and H344Y were neutral, implying that the aromatic pi system of H344 may play a role at this position. Exchanges at D315, which may form a hydrogen bond to K346 to stabilize loop 6, were beneficial in cases where a hydrogen bond could be retained (D315E, D315S).

Residues in a stretch between G397 and I411 exhibited a consistently strong negative effect upon substitution under all conditions. This stretch forms an α-helix positioned above an opening that connects the cavity where RuBP binds to the protein surface (Supp. Fig. 5A, 5B).

The very low tolerance to mutations in these residues likely reflects the importance of maintaining this structural opening. Loss or alteration of this tunnel could prevent the enzyme from properly binding or releasing RuBP, 3PGA, and 2PG.

CbbM contains a nine residue insert (positions 78-87) that is not present in well-investigated form II rubiscos such as those from *R. rubrum* and *Gallionella GW1SB* or in form I rubiscos from spinach and cyanobacteria. Structural predictions of the entire dimeric protein using both homology modeling and AlphaFold showed that this insert is in a loop region. Mutations at V84 and K85 within this insert generally increased fitness. Other positions at which, on average, replacements had a positive effect were Q128 and K219, which are part of a beta-sheet in the N-terminal domain and an alpha helix in the C-terminal domain, respectively.

Residue W114 is at the dimer-dimer interface, an area presumed to be less tolerant to mutation. However, mutations at W114 are among the highest fitness variants in the saturation library, and the effect was stronger in the aerobic than in the anaerobic condition. Tunnel analysis using CAVER 3.0 revealed that residue W114 lies adjacent to a continuous cavity or tunnel that extends through the enzyme and connects the two active sites of the dimer (Supp. Fig. 5C).

### Combinatorial mutagenesis reveals extensive epistatic interactions in rubisco

The average fitness of CbbM decreased with the number of introduced mutations (Fig. 2B), as reported for other proteins (48). However, we found several higher-order variants with fitness exceeding that of single-site variants. Nevertheless, the extent of such fitness increase was not high, suggesting that the zero-shot predictions for multi-site exchanges are significantly hindered by unseen epistatic effects. An illustrative example of epistasis is the three-point variant (H127V, K247E, S432A), and the single-point variant H251A, which both showed negative fitness compared to CbbM_base. However, the combined four-point variant showed significantly higher fitness (Fig. 2D).

For each of the 21 exchanges in the combinatorial library, we calculated the additive fitness of adding that exchange to a variant. Most exchanges could be either beneficial or detrimental, though there were some exceptions. Exchanges at V323 (V323A and V323S) were detrimental in all variants (Fig. 2C). The residue V323 appears to contact an extended tunnel which connects the protein surface to the active sites (Supp. Fig. 5C and 5D), and the negative impact of substitutions suggests that this tunnel is essential for catalysis in CbbM_base. By contrast, mutations at H127, Q392, and S432 were rarely detrimental and frequently beneficial. Interestingly, these residues are on the surface of the CbbM.

To validate results from the high-throughput fitness screen, we cloned several CbbM variants into the *Synechocystis* rubisco-depleted strain and measured growth rates in batch cultivations. This validation included variants with high fitness values from the saturation mutagenesis library; variants K85N, F90A, and Y319T and combinatorial library; C4 (H127V, K247E, H251A, S432A), C5 (H127V, K247A, H251K, Q392E, S432I) and C6 (H127V, M140A, K247A, H251A, Q392E, S432T). All of the resultant *Synechocystis* clones showed faster growth than CbbM_base as well as CbbM_wt clones, in agreement with fitness data from library screening (Fig. 2E, 2F). To determine whether the improved fitness was due to altered kinetic parameters or variant solubility (60), we compared CbbM abundances with western blotting (Supp. Fig. 6). In all cases, differences in CbbM expression were minor. However, the western blots do not account for the activation state of CbbM variants. A putative Rubisco activase is present in the *Gallionella* genome, but was not included in this study.

### Zero-shot models predict fitness of single amino acid exchange mutants but not multi-site mutants

The CbbM fitness data allowed us to test the ability of the zero-shot models EVmutation, DeepSequence and an MSA Transformer ensemble (44,45,47,61) to predict fitness scores based on protein sequence. These models use phylogenetic data as captured by MSAs to predict the effect of amino acid exchanges. EVmutation is a statistical model capturing site-specific amino acid biases as well as co-dependencies in each pair of sites. DeepSequence and MSA Transformer are neural networks based on a variational autoencoder and a transformer architecture, respectively. Fitness predictions from all three models were highly correlated (Supp Fig. 7), and all performed well in predicting the effect of single amino exchanges (r=0.38-0.45), in line with their reported results on benchmarking datasets (49) (Fig. 3A, Supp. Fig. 8). However, predictions of all models correlated negatively with experimentally determined fitness of higher order variants (Fig. 3A, Supp. Fig. 8). The inclusion of exchanges V323A and V323S in many combinatorial mutants, which was universally negative despite predictions, has an outsize effect on correlation to combinatorial library data. Removal of variants containing V323 mutations improved fitness predictions (Pearson’s r increased from -0.30 to +0.24 for EV mutation; (Supp. Fig. 9)).

### Using labeled fitness data to guide rubisco engineering

To assess whether the CbbM fitness dataset could inform a second round of CbbM mutagenesis, we tested both naive and machine-learning guided protein engineering approaches. We first constructed a simple additive model that assumes independence and additivity of fitness effects from single-site exchange variants. This additive model predicted the fitness scores of the library’s higher-order variants well (Fig. 3B, 3C, Supp. Fig. 10, Supp. Data file 1). Inspired by this model, we tested whether combining positive exchanges would further increase fitness. We created six additive CbbM_base variants where we combined single-site exchanges: S2 (V84R, K85N), S3 (V84D, K219I, D315T), S4 (F90C, W114K, V256L, M331I), S8 (V84D, F90C, W114K, Q128D, K219I, V256L, M331I, H344S), and two variants where we added positive single-site exchanges to high-fitness combinatorial variants: Ad1 (H127V, K247A, H251A, Q392E, S432T, and W114I) and Ad2 (H127V, K247A, H251A, Q392E, S432T, W114I and K346I). We also trained ProteinNPT on the entire experimental fitness data set. ProteinNPT is a non-parameteric transformer that learns from experimental data connected to CbbM sequences as well as MSA Transformer predictions of the same sequences (62,63). After training on single- exchange variants, ProteinNPT could predict the fitness of the combinatorial mutants well (Supp. Fig. 10). From 200 variants predicted by ProteinNPT to have high fitness scores, we selected 11 (ML1-ML11) to test experimentally. These variants contained between 6 and 12 mutations compared to CbbM_base (Supp. Table 6). We then transformed the rubisco-depleted *Synechocystis* strain with all second-generation variants and cultivated the strains individually.

Additive variants S2 and Ad1 showed significantly higher growth than CbbM_base and CbbM_wt, and variant Ad1 exceeded growth of all other variants in this study (Fig. 3D).

Furthermore, the Ad1 variant exhibited enhanced growth in 1% CO₂ and ambient air conditions compared to CbbM_base and other first-generation variants (Supp. Fig. 11). Among ProteinNPT variants, two showed higher growth than CbbM_base, ML4 and ML7, each with six substitutions. However, most *Synechocystis* strains containing second-generation CbbM variants did not grow better than *Synechocystis* containing CbbM_base. Of the additive variants, three of six showed growth similar to a negative control lacking a functional rubisco (S3, S4, S8), and of the ML variants 8 of 11 did not impart growth. Thus, neither the additive model nor the ProteinNPT- assisted model had high accuracy in extrapolating fitness values outside the range present in the dataset. Here again epistasis complicated protein engineering. While variant Ad2 differed from variant Ad1 in only a single exchange which independently was beneficial (K346I), this variant had severely impaired growth. To assess whether adaptive mutations were transferable to other genetic backgrounds, we created a composite mutant (“Ad1*”) incorporating all substitutions from Ad1 but excluding the base variant mutations (S162P and G442R). This variant failed to grow under elevated CO₂ conditions, indicating that the Ad1 exchanges are specifically beneficial for the CbbM_base starting variant and not CbbM_wt.

### Catalytic parameters of higher-order rubisco variants

We next determined the catalytic parameters of C6, the highest fitness zero-shot variant, and Ad1, the highest fitness 2^nd^ generation variant, as well as CbbM_base, and CbbM_wt using an *in vitro* ^14^CO2 assay (64–66). The variants were expressed in *E. coli* and purified without removing the C-terminal polyhistidine tag, which was previously reported to not affect enzyme activity (67). Kinetic parameters for CbbM_wt were published previously, and our results were similar (21). CbbM_base exhibited a 30% lower *k*cat^C^ and 55% higher *k*cat^O^ compared to CbbM_wt. Thus, the starting point for the library had impaired kinetics compared to CbbM, which was not apparent in our previous fitness screen, presumably because expression of CbbM variants in that study was from an episomal plasmid with a strong promoter (41). Both C6 and Ad1, which imparted better growth in *Synechocystis* cultures than CbbM_base, had markedly lower *k*cat^O^ and subsequently lower *k*cat^O^/KO (Fig. 4C). Additionally, Ad1 had a significantly increased *S*C/O compared to CbbM_base and C6 (Fig 4A). Ad1 contains the exchange W114I, which may affect gas tunnels (Fig. 4J-L). C6 had a 25% higher *k*cat^C^ than CbbM_base , and restored turnover number to that of CbbM_wt, though Kc was also increased. The exchange M140A distinguishes C6 from Ad1, and alanine in this position is present in rubiscos from *R. rubrum*, *S. oleracea,* and *Synechocystis* sp. PCC 6803 (Supp. Fig 1). A result of these changes in individual parameters is that *k*cat^C^/KC and *k*cat^O^/KO were correlated among all variants (Figure 4G- I), while *k*cat^C^ and *k*cat^O^ were not correlated.

We also tested thermal stability and oligomerisation of selected CbbM variants. We found that variants displaying increased fitness in aerobic growth generally had a lower melting temperature as measured by NanoDSF (Supp. Fig. 11). This reduced stability may reflect an altered protein-protein affinity at the dimer interface causing a reduction in protein rigidity. One such example could be the exchange W114I, which may open up cavities in between the dimers (Supp. Fig. 5) To see if the mutations had any effect on the quaternary structure (24), we performed Mass photometry and compared Ad1 to CbbM_base. Both variants were found to be present as dimers in 95 - 97 % of the population.

## Discussion

Here we report fitness data on more than 15,000 mutants of the form II *Gallionella* rubisco. When taken together with other high-throughput, labeled fitness screens (29,68), this dataset can form the basis for training rubisco models. As an alternative to full-protein DMS and error-prone PCR methods, we employed zero-shot predictors based on a rubisco MSA containing 10,492 sequences (Meff = 2,438). We reasoned that this approach was more likely to result in rubiscos of higher fitness. It is noteworthy that, even though our zero-shot library covered just a fraction of CbbM residues (67/470, 14%), approximately 8% of zero-shot variants showed an increased fitness compared to CbbM_base. This high number could reflect a general advantage of zero-shot predictors, or simply that there is less selective pressure on oxygenation in *Gallionella* rubisco, so that more oxygen insensitive mutants are “nearby,” in sequence space. As the MSA used to train the zero-shot predictors contains thousands of rubiscos that are likely more resistant to oxygen than CbbM, the fitness predictions may be expected to tend toward oxygen resistance. Comparison to observed fitness showed that the zero-shot predictors were accurate in predicting fitness effects of single-site mutants, but less accurate at predicting the fitness of multi-site variants. A large factor was the presence of a single epistatic exchange in the multi-site library, though when this was accounted for prediction was still only moderate. Poor prediction of epistasis could indicate a bias in the underlying MSA used by these models, where diversity of sequence is more critical than the MSA depth (61). To mitigate this problem, emerging zero-shot predictors for proteins enhance accuracy by incorporating protein structures as well as MSAs (69,70).

We used our labeled fitness dataset to test two approaches for enhancing fitness of CbbM_base. We compared a directed evolution-like recombination strategy, in which beneficial mutations are combined, and a machine-learning approach, where the partial fitness landscape is learned and used to predict fitness of new variants. Previously, it was estimated that machine-learning guided directed evolution marginally outperforms a recombination approach, in that mutants with optimal fitness could be achieved with fewer rounds and screened variants (71). In our study, the fraction of 2^nd^ generation variants that were functional and higher fitness than CbbM_base were similar between the two methods (approximately 50% and 15%, respectively). It is likely that the CbbM fitness could be further enhanced by additional rounds of validation, generation and sampling, using emerging ML approaches (38,72).

Our results give insight into structure-function relationships in rubisco. Replacements that affect the flexibility of loop 6 affected fitness, with stabilizing mutations having a positive effect, as has been reported previously for other rubiscos (28,73,74). Fitness-enhancing mutations were often found at the dimer interface. Interpretation of fitness data with CAVER analysis suggests that substrate access tunnels may be critical for gas entry and exit. Mutations at site W114 that destabilise the dimer interface, as indicated by a lower Tm, while simultaneously opening a tunnel. Recently, it was shown that minimal mutations to the hexameric RbcL6 from *Gallionella GWS1B* could shift the quaternary structure to an L2 dimer, which also significantly increased the apparent O2 binding constant (24). Furthermore, a directed evolution study of the RbcL6 from *Gallionella GWS1B* found that multiple mutations in the dimer interface affected oxygen sensitivity of the enzyme (22).

The cyanobacteria screening system highlights challenges in using *in vivo* growth rates to guide optimization of specific catalytic parameters. In our previous work, we identified CbbM_base as having similar fitness to CbbM_wt. In this study, where all CbbM variants were expressed from a weaker promoter, there was a small but noticeable reduced cell fitness of CbbM_base compared to CbbM_wt (Fig. 2E). Therefore, it is important for growth-coupled fitness screens that total enzyme activity is not too high as to mask changes in kinetic parameters. In the case of rubisco, efforts to increase carboxylation rate often result in mutants that are more highly expressed instead, particularly in Form I rubiscos (17). This did not appear to be the case in this study, as western-blotting showed similar or even reduced expression of rubisco variants that enhanced growth. There was a clear trend that *in vivo* fitness was more enhanced in the 20% O2 condition compared to 0% O2 condition, which suggests that increased growth rate imparted by rubiscos in cyanobacteria is due to the reducing oxygenation reaction. The determined kinetic parameters supported this, as both variants C6 and Ad1, which showed significantly higher growth than CbbM_base and CbbM_wt, displayed lower catalytic efficiencies for the oxygenation reaction *k*cat^O^/KO but also lower carboxylation efficiency *k*cat^C^/KC and *k*cat^C^/KC^air^. The results are also consistent with a recent directed evolution study of the RbcL6 from *Gallionella GWS1B,* which showed that the majority of mutants imparting improved growth of RDE *E. coli* grown in air had reduced oxygenation rates (22). We used a relatively high CO2 gas feed of 5% vol/vol; growth of CbbM mutants at lower CO2 concentrations was extremely low (Supp. Fig. 10). We assumed that this high substrate would have exceeded KC and put selection pressure on *k*cat^C^. In the case of variant C6, *k*cat^C^ was increased by 30% relative to CbbM_base. These results show it is possible to alter *k*cat^C^ and *k*cat^O^ independently in rubisco, parameters which describe the hydration and cleavage and release of carboxyketone and peroxiketone intermediates, respectively (8).

However, changes in *k*cat/KM were correlated for oxygenation and carboxylation reactions. Although we characterised only a few variants, this result is consistent with a model that these parameters, which describe gas addition, are mechanistically coupled (8).

Enhanced rubiscos have implications for bioproduction. In metabolic engineering applications with CO2-fixing bacteria, where product synthesis may be limited by rubisco activity (5), fast rubiscos with poor CO2 affinity may be good candidates for incorporation into carboxysomes, or cultivated at high CO2 conditions to exploit high *k*cat. Fast rubiscos optimized for activity in oxygenic photosynthetic cyanobacteria, such as through significantly reduced oxygenation, may be more readily transferred to plants due to their closer evolutionary relationship. However, this application area is more complex, as the extent of limitation from rubisco kinetics is highly species specific. Recently, it was shown that the form II rubisco from *R. rubrum,* which has a significantly lower *k*cat than the rubiscos reported here, was sufficient to sustain potato growth similar to wild type at elevated CO2, suggesting enzyme turnover is not limiting (75). In conclusion, our extensive fitness dataset could form the basis for continued engineering of rubisco to obtain both fast and selective variants.

## Materials and Methods

### Experimental design

The objective of this study was to identify variants of *Gallionella* rubisco, CbbM, which increased the growth rate of a cyanobacterial host strain, *Synechocystis*, when replacing endogenous *Synechocystis* rubisco and compared to a starting CbbM variant. We performed controlled laboratory experiments including the pooled screening of a library of mutant variants at different growth conditions that should select for different properties of the screened enzyme. In a next step, promising variants were characterized further both *in vivo* as well as *in vitro*.

### Library design, zero-shot predictions and ProteinNPT

The screened library consisted of a saturational library, in which single amino acid exchanges were introduced into each separate variant, and a combinatorial library, into which up to seven amino acid exchanges were introduced at the same time. For the saturational/DMS part of the library, we predicted the fitness of Rubisco variants with a single amino acid exchange using the EVmutation (44,45) web server (https://v2.evcouplings.org/) and DeepSequence (47). We used the amino acid sequence of the N-terminally Strep-tagged CbbM_base variant (GenBank identifier OGS68397.1 with amino acid exchanges A230E and G422R) as input for the EVmutation web server (https://v2.evcouplings.org/) with default settings (bitscores 0.1, 0.3, 0.5, 0.7; 5 search iterations; sequence database UniRef90; position filter 70%; sequence fragment filter 50%; removing similar sequences 90%; downweighting similar sequences 80%; statistical inference model: pseudolikelihood maximization). The web server identified a bitscore of 0.3 as optimal for alignments, which led to a multiple sequence alignment of 10,492 sequences (number of effective sequences Meff = 2,438; number of redundancy-reduced identified homologs divided by confidently aligned positions in the alignment Neff/L = 6.5). This alignment was used for all subsequent tools which relied on a MSA. To run DeepSequence, we adapted code by Hsu et al., (48) for training the DeepSequence model, used a conda environment provided by Wittmann et al., (37) and adjusted the DeepSequence code slightly. We picked the subset of residue positions for which the EVmutation epistatic score was above 1.0 or which were among the top 100 predictions of DeepSequence and a EVmutation epistatic score above -1.0. In addition to EVmutation/DeepSequence zero-shot predictions, part of the library consisted of lysine residues in close proximity to residues that promote chemical acetylation (Supp. Table 1). These were chosen based on the assumption that Rubisco acetylation at a lysine residue might have a regulatory effect (51).

Exchanges to be included in the combinatorial library were chosen as described previously on the basis of “*in silico* evolution” using either EVmutation or DeepSequence as a fitness predictor. For EVmutation, we used 1000 trajectories of *in silico* evolution with 500 steps for a first step, in which all possible exchanges could be introduced and a second step, in which only a subset of 20 residues, which occurred most frequently during the first step, could be exchanged.

We ran DeepSequence predictions on a Google Cloud Product n1-highmem-2 virtual machine with an NVIDIA T4 GPU. When using DeepSequence as fitness predictor, we performed 100 trajectories of *in silico* evolution for 500 steps with all possible amino acid exchanges. This was followed by another 100 trajectories of *in silico* evolution for 200 steps after limiting to the same subset of most frequently occurring 20 exchanges as mentioned above. All trajectories were run at a temperature of 0.03. The most frequently occurring amino acid exchanges predicted by the latter rounds of *in silico* evolution were included in the combinatorial library.

Zero-shot predictions for the entire library were performed using an ensemble of the MSA Transformer model based on five different seeds using code provided by (https://github.com/OATML-Markslab/ProteinNPT/) on an AWS EC2 g6e.xlarge instance with an NVIDIA L40S Tensor Core GPU. When fitness values were available, ProteinNPT was, in a first step, trained on fitness data of the complete saturational part of the library in the multi-objective setting. This training data set included fitness values for the single amino acid exchange variants underlying the combinatorial library. The trained model was used for predicting combinatorial variants for which normed fitness values were available. In a second step, ProteinNPT was trained on the whole library and the resulting model was used to predict the fitness of variants which seemed to be promising based on the fitness effect of the separate exchanges. These numbers were compared with the naive additive model and based on both, variants to be tested experimentally were picked. For training of ProteinNPT, the same AWS EC2 instance as mentioned above was used.

### Library synthesis and plasmid preparation

Libraries were synthesized by GenScript (Amsterdam, Netherlands). After receiving the plasmid pools, libraries were transformed via electroporation into CopyCutter™ EPI400™ *E. coli* cells (Lucigen, epicentre) and colonies corresponding to a 70X - 100X coverage of each library were pooled in LB medium separately for the combinatorial and the saturational library. Aliquots were frozen in 25% (v/v) glycerol, 0.5X LB and aliquots were kept at -80°C until further usage.

For plasmid preparation, one aliquot per library was thawed in 50 ml LB medium containing kanamycin (50 µg/ml) and grown overnight at 37°C. The next day, cultures were diluted 1:100 in LB containing 1X CopyCutter™ solution (Lucigen, epicentre). After 4 hours of growth in baffled Erlenmeyer flasks (37°C), cells were harvested and plasmid prepared using MiniPrep kits (Thermo Scientific).

### PacBio sequencing

To assign the N20 barcodes to mutant variants, plasmid DNA extracted from the *E. coli* mutant library was sequenced. To do so, induction of a high plasmid copy number of the pool of CopyCutter EPI400 *E. coli* cells was performed according to the manufacturer’s protocol.

Plasmid was extracted using a GeneJET Plasmid Miniprep Kit (Thermo Scientific™) and DNA concentration was measured using a Qubit™ 1X dsDNA High Sensitivity (HS) assay kit (Invitrogen™). Saturational and combinatorial plasmid pools were linearized in separate reactions of three to five µg plasmid DNA with single-cutter FastDigest restriction enzymes EcoRI, SmaI or Alw44I (Thermo Scientific™). Linearized plasmid was extracted from agarose gels using the Zymoclean Gel DNA Recovery Kit (Zymo Research). Libraries were pooled according to their theoretical number of members and the pool was purified using AMPure XP beads (Beckman Coulter) with a protocol adapted for large DNA fragments. One volume of AMPure XP beads were mixed with the sample and incubated at room temperature for 10 minutes. Beads were separated on a magnetic rack for 2 minutes and supernatant withdrawn. Beads were washed twice with freshly prepared 80% (v/v) ethanol and incubation periods of 1 minute. After a short spin, remaining ethanol was withdrawn and beads were air dried for 2 minutes. DNA was eluted by adding 0.5 X H2O, incubating it for 10 min with the beads, removing beads from the eluate by keeping the tube on the magnetic rack for 2 minutes and then removing the supernatant. This eluate was handed to the national genomics infrastructure (NGI) Sweden for further library preparation. Adapters were added using the SMRTbell prepkit 3.0. The library was sequenced using a Sequel Sequencing Kit 2.0 and a Sequel™ SMRT® Cell 8M v3 Tray on a Sequel II machine.

### Bioinformatic Analyses of PacBio data

PacBio sequences were mapped using minimap2 v2.28 (r1209) (76) and its CCS-specific preset to a 1,675 bp long template spanning the promoter region (P*trc*), the coding sequence of *cbbM* and a small region up- and downstream of the N20 barcode. The resulting sam-file was filtered for aligned reads with a mapping quality above 5 (MAPQ > 5) using the samtools v1.13 (77) view command. Further processing was mainly performed using a Jupyter notebook for Python 3.9.7 (PacBio barcode processing.ipynb). First, aligned reads were extracted from the resulting sam file and converted into fasta format. To extract coding sequences and barcodes from this file, cutadapt v3.5 (78) was run twice using two different sets of paired adapters corresponding to the sequences exactly up- and downstream of the *cbbM* coding sequence and the N20 barcode, respectively. Untrimmed reads and barcode reads shorter than 15 nt were discarded. Coding sequences were assigned to barcodes according to the PacBio read identifier and coding sequences were translated to protein sequences. Protein products shorter than 300 amino acids were discarded. For barcodes occurring more than once, the assigned coding sequences were compared and the sequence occurring most frequently was used as consensus sequence. In case there were ties, these barcodes were discarded. All remaining barcodes were combined into one file, amino acid exchanges in comparison to the CbbM sequence that was underlying the library (CbbM with A230E and G422R) were determined and the combination of barcodes and assigned mutations were formatted to be compatible with downstream applications. In total, 235,598 barcodes were identified. Of these, 205,150 barcodes were assigned to 17,839 mutant variants expected based on the initial library design. We expected 17,983 mutant variants from the initial library design – 1,805 variants belonging to the saturational part, 16,200 to the combinatorial, which are partially represented in the saturational part. The remaining barcodes represented mutant variants longer than 300 amino acids covered by more than one barcode. All variants expected on base of the library design and missing in the *E. coli* library belonged to the combinatorial library.

Illumina sequencing data from the library cultivation at continuous light and a gas feed of 5% CO2, 95% N2, 0% O2 were mapped to the resulting barcode library. For further analyses, barcodes were only kept if more than 30 reads from these 30 5% CO2, N2 samples mapped to their sequence. The resulting barcode library consisted of 32,549 barcodes, which represented 14,329 expected mutant variants. Only the 30 barcodes with the highest counts assigned to the library base variant, CbbM with A230E and G422R, were kept in this library. In total, 8,968 mutant variants were represented by more than one barcode.

### Strain construction Synechocystis

All *Synechocystis* strains and annotated sequences of plasmids used in this study are provided on GitHub repository. All library variants were transferred into the receiver *Synechocystis* strains with an inducible CRISPRi suppression system targeting the native rubisco as described in Hoffman et al., 2025 (41) by natural transformation. The transformation with the library yielded 60,000 colonies. The combinatorial and saturation part of the library was combined in a ratio of 10:1. The combined library was mixed in 25% (v/v) glycerol, 0.5x BG-11 (HEPES), snap-frozen, and stored at -80 °C until further usage.

All variants to be validated and other second-generation variants were ordered as a gene block and cloned into the pEERM-based library plasmid in XL1 blue E. coli. The variants were transformed into the CRISPRi strain via natural transformation and individual colonies were picked and confirmed via cPCR.

### Pooled growth of *Synechocystis* library

The pooled cultivations was performed in a 8-tube Multi-Cultivator MC-1000-OD bioreactors (Photon System Instruments, Drasov, CZ) using the pycultivator-legacy package (https://gitlab.com/mmp-uva/pycultivator-legacy) (79) and as described (80) with the following modifications: Light intensity was kept at 300 µmol photons m-2 s-1. The turbidity threshold was set to OD720nm=0.2, if the measurement was exceeded for two measurements in a row, automatic dilution occurred. The library was cultivated in 8 separate growth channels, so 8 replicates per condition, inoculated from the same pool. After the pooled library had acclimated and reached stable cultivation with regular back-dilutions, the native rubisco was suppressed by induction of the CRISPRi system by adding aTc (2 µg ml-1) to cultivation tubes and feeding bottles. Addition of aTc marked generation 0. Growth data was analyzed using the ShinyMC web application (https://github.com/m-jahn/ShinyMC, v0.1.1) growth rates and doubling time was calculated as mentioned in Hoffmann et al., 2025 (41). Cells were harvested after approximately the 4th, 8th, and 10th generation by centrifuging 12 ml culture (4,500 x g, 4°C, 10 min).

Supernatant was removed completely and cell pellets were frozen at -20°C until further usage for NGS preparation. For continuous light, oxygen-free gas feed, cultivation in one of the channels stopped after the fourth generation due to technical difficulties.

### Library preparation and next-generation sequencing

The NGS library preparation was done as described in Hoffmann et al., 2025 (41) with the following changes: DNA was extracted using a GeneJET Genomic DNA Purification Kit (Thermo scientific) following the protocol for Gram positive bacteria and eluted in 40 µl deionised water. After preparation, the library was run on a NextSeq 2000 system (Illumina) using NextSeq 1000/2000 P2 reagents v3 (Illumina) according to the manufacturer’s instructions.

### Illumina NGS bioinformatic analysis and exploratory data analysis

Sequencing data was analyzed using the Nextflow nf-core-crispriscreen pipeline (Miao & Jahn et al., 2023) (commit d9e9b1812b2e093a49f290cd0b04fc407ff1d5a2 from 31st Oct 2024) with the following input deviating from the defaults: the sequence GTCTAGAatcgccgaaagtaattcaactccattaa…TCTAGATGCTTACTAGTTACCGCGGCCA was used for the parameter --five_prime_adapter, the error rate was set to 0.2, filter_mapq was set to 1 and both run_mageck as well as gene_fitness were set to false. Briefly, the pipeline performs read trimming with cutadapt (78), maps reads with bowtie2 (81), data normalization with DESeq2(82) and calculation of fitness scores for individual barcodes. The fitness of a specific barcode at a given condition was determined as the area under the curve given by the log2FC over time *t* normalized by total cultivation time given in generations as given by (80):

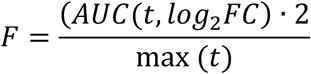

Custom R Markdown and python scripts were used for subsequent analyses. Fitness values of barcodes associated with a specific mutant variant were compiled to a weighted mean fitness value *Fwmean* per mutant variant and condition by weighting according to the Pearson correlation of each specific barcode, *ri*, with the rest of the barcodes associated with this variant, and the inverse of the maximum standard error on the log2FC values given for a specific barcode by DESeq2:

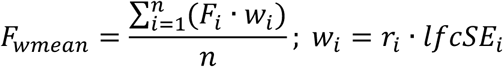

Adjusted p values were calculated by comparing the fitness values of the set of barcodes associated with a specific protein variant and the barcodes associated with the protein variant underlying the complete library (CbbM with S162P, G422R) using the two-sided Wilcoxon signed rank test and Benjamini-Hochberg correction (significance level 0.1).

For plotting, weighted mean log2FC were compiled from separate log2FC values of different barcodes by following the same strategy as described above, i.e. by weighting by the weighting factor *wi* given by the Pearson correlation of a given barcode and its maximum standard error on the log2FC.

We adapted the methodology of Miton and Tokuriki (83) to analyze the effect of exchanges on intermediate variants. They compared the fitness effect of an amino acid exchange *i* on the starting enzyme background, which would be in our case CbbM_base, and on higher-order, intermediate variants. To do so, they calculated the ratios of the fitness of the variants containing the exchange compared to the base variant fitness (Fbase,i=Fbase+iFbase) and a higher-order, intermediate variant *j* (Fj,i=Fj+iFj). We adjusted their approach as follows: Fitness values below 0.1 were set to 0.1 as we considered them as catalytically inactive, based on the distribution of normed fitness values of K200 variants. Deviating from Miton and Tokuriki (83), we considered an exchange with 0.8<Fj,i<1.25 as neutral, one with Fj,i<0.8 as detrimental and with Fj,i>1.25 as beneficial. We adapted these cut-offs on the basis of the value distribution in our data set.

### Further data analyses

Multiple sequence alignments were created using Clustal Omega (84,85) and visualized with JalView v2.11.1.4 (86). Protein structures were visualized using UCSF ChimeraX v1.8 (87) and PyMol v2.5.0 (88). The solvent-accessible surface area was calculated using the ChimeraX command “measure sasa” and relative SASA using theoretical MaxASA values by Tien et al., 2013 (89).

### Proteomics mass spectrometry sample preparation and analysis

Cell pellets were collected as described for photobioreactor batch cultivations. They were lysed by bead beating (FastPrep-24 5G lysis machine, MP Biomedicals). Proteomics analysis was performed on a Q-exactive HF Hybrid Quadrupole-Orbitrap Mass Spectrometer coupled with an UltiMate 3000 RSLCnano System with an EASY-Spray ion source. A more detailed protocol can be found described in Hoffmann et al., 2025 (41).

### Western Blot and Nano differential scanning fluorimetry (nanoDSF) Analysis

Solubility and stability comparison between mutant variants were performed as described in Hoffmann et al., 2025 (41).

### *In vitro* kinetic characterization

In preparation for in vitro verification the gene for a select number of variants were cloned into pET28a-6xHis-KmR and transformed into *E. coli* BL21(DE3). For each CbbM variant, a single colony was used to inoculate 20 mL of LB (50 µg/ml Km), and incubated overnight at 37 °C. The following day, 200 mL 2YT (50 µg/ml Km) culture were inoculated to an OD of OD600 nm of 0.1. 0.5 mM isopropyl-β-d-thiogalactopyranosid (IPTG) was added to induce protein expression when an OD600 nm of 0.6-0.8 was reached, cultures were moved to 16 °C and incubated for 16 h before harvested by centrifugation (3005g, 4 °C, 20 min). The harvested cells were lysed using B-PER and purified using Ni-NTA agarose beads (Qiagen) as described in Hoffmann et al., 2025 (41) The proteins were put in a buffer with 18 mM MgCl2 and 18 mM EPPS, and then bicarbonate was added to 50 mM. The resulting purified proteins were immediately frozen in liquid nitrogen and stored at −80 °C until measurement.

Extracts were supplemented with carrier-free NaH^14^CO3 to adjust the specific radioactivity to 3.7 x 10^10^ Bq mol^-1^ and preactivated at 25°C for 20 min. Rates of Rubisco ^14^CO2- fixation were measured at 25°C in 7 ml septum-capped glass scintillation vials containing 400 µl of Assay Buffer (100 mM Bicine pH 8.1, 20 mM MgCl2) and ∼ 100 W-A units of carbonic anhydrase (C3934 Merck, USA), bubbled either with 100% N2 gas or CO2- free synthetic air (21% O2, 79% N2) for 1 h. After that, one of the eight different concentrations of ^14^CO2 ranging from 16 to 500 µM and 1.6 mM of RuBP (synthesized and purified as explained in Kane et al. (1994) (65) were added. Assays were started by the injection of 20 µl of pre-activated extract and quenched after 1 min by the addition of 0.2 ml of 10 M formic acid. After removal of acid-labile ^14^C by evaporation, acid-stable ^14^C was determined by liquid scintillation counting (Tri-Carb, Revvity USA). The Michaelis–Menten constant for CO2 (*K*c) and the maximum carboxylase activity (*V*_max_ ^c^) was calculated as in Capó et al. (64). The Michaelis–Menten constant for O (*K* ) were calculated in each biological replicate by a linear fit of the *K*c measurements obtained under 0% O2 and under 21 % O2. The carboxylation turnover rate (*k*cat^c^) was calculated by dividing *V*_max_ ^c^ by the concentration of Rubisco active sites, which was quantified from the same protein extracts by 2ʹ-carboxyarabinitol-1,5-bisphosphate (^14^C-CABP, 130 µM final concentration) binding (Kubien et al., 2011) (66). Concentrations of CO2 in solution were calculated assuming an acid dissociation constant for carbonic acid (pKa1) of 6.11 and a solubility constant for CO2 of 0.034 mol l^−1^ atm^−1^ at 25°C and using accurate measures of the pH (NBS scale).

For *S*C/O, CO2-free protein extracts were obtained as explained before, using a Desalting Buffer without NaHCO3 and bubbled with 100% N2 gas. *S*C/O was analyzed by using [1-^3^H] RuBP, following the method of Kane et al. (1994) (65) with the modifications of Capó et al. (2020) (64). The assay was equilibrated with a gas mixture of 99.95 % O2 and 0.05% CO2, and *S*C/O was calculated using a CO2/O2 solubility ratio of 0.038 at 25°C.

### Protein model prediction and Tunnel analysis

The three dimensional structure of the wild-type CbbM was initially predicted using homology modelling using YASARA Structure (v. [-]). The protein sequence was submitted to the YASARA Homology Modeling module, which identified suitable structural templates through BLAST and PSI-BLAST searches against the Protein Data Bank. Templates were selected based on sequence identity, alignment quality, and structural resolution (see Supplementary info). Each model was refined using YASARA’s integrated optimization and energy minimization pipeline with the AMBER03 force field in explicit solvent. Resulting models were ranked by YASARA’s Z-score, and the top-scoring model was chosen.

The structure of the designed variants used for tunnel analysis were predicted using the AlphaFold Protein Structure Database prediction server (DeepMind/EMBL-EBI) (90). The protein sequence, without streptag, was feed to the server and the presence of magnesium ion was incorporate. The “Multimer” option was selected to enable prediction of the dimeric assembly.

Two identical copies of the sequence were provided as input to generate a homodimer model. The server automatically generated multiple sequence alignments using its standard suite of sequence databases and executed the AlphaFold-Multimer prediction pipeline. For each submission, five structural models were produced. Confidence was evaluated using: pLDDT for per-residue accuracy; ipTM and pTM scores for global and inter-chain prediction confidence; Predicted Aligned Error (PAE) maps to assess chain–chain orientation reliability. The highest-ranked model according to AlphaFold’s internal ranking metric was selected for downstream analysis.

The predicted structures were then analysed using Caver Web 2.0 (91), using residue K200 as a starting point and with the following parameter inputs: probe_radius 0.9, shell_depth 4.0, shell_radius 3.0, clustering_threshold 3.5, max_distance 3.0, desired_radius 5.0.

## Acknowledgments

We thank Michele Russo (KTH) for assistance in sample preparation and bioreactor cultivation, Sara Lupacchini (KTH) for assistance in bioreactor cultivation, and Cenk Gurdap (Karolinska Institutet) for assistance with mass photometry.

## Author contributions

Conceptualization: EPH, UAH, AK, RM, KS Methodology: EPH, UAH, AK, RM, AZS Investigation: EPH, UAH, AK, AZS, ES Visualization: UAH, AK

Supervision: EPH, UAH, AK Writing—original draft: EPH, UAH, AK

Writing—review & editing: EPH, UAH, AK, AZS

## References

1. Croce R, Carmo-Silva E, Cho YB, Ermakova M, Harbinson J, Lawson T, et al. Perspectives on improving photosynthesis to increase crop yield. Plant Cell. 2024 Oct 1;36(10):3944–73.

2. Carmo-Silva E, Sharwood RE. Rubisco and its regulation—major advances to improve carbon assimilation and productivity. J Exp Bot. 2023 Jan 11;74(2):507–9.

3. Zhu XG, Portis Jr AR, Long SP. Would transformation of C3 crop plants with foreign Rubisco increase productivity? A computational analysis extrapolating from kinetic properties to canopy photosynthesis. Plant Cell Environ. 2004;27(2):155–65.

4. Flügel F, Timm S, Arrivault S, Florian A, Stitt M, Fernie AR, et al. The Photorespiratory Metabolite 2-Phosphoglycolate Regulates Photosynthesis and Starch Accumulation in Arabidopsis. Plant Cell. 2017 Oct 1;29(10):2537–51.

5. Hudson EP. The Calvin Benson cycle in bacteria: New insights from systems biology. Calvin Benson Bassham Cycle. 2024 Mar 1;155:71–83.

6. Wilson RH, Thieulin-Pardo G, Hartl FU, Hayer-Hartl M. Improved recombinant expression and purification of functional plant Rubisco. FEBS Lett. 2019;593(6):611–21.

7. Prywes N, Phillips NR, Tuck OT, Valentin-Alvarado LE, Savage DF. Rubisco Function, Evolution, and Engineering. Annu Rev Biochem. 2023 June 20;92(Volume 92, 2023):385–410.

8. Flamholz AI, Prywes N, Moran U, Davidi D, Bar-On YM, Oltrogge LM, et al. Revisiting Trade-offs between Rubisco Kinetic Parameters. Biochemistry. 2019 Aug 6;58(31):3365– 76.

9. Tcherkez GGB, Farquhar GD, Andrews TJ. Despite slow catalysis and confused substrate specificity, all ribulose bisphosphate carboxylases may be nearly perfectly optimized. Proc Natl Acad Sci. 2006 May 9;103(19):7246–51.

10. Bouvier JW, Emms DM, Kelly S. Rubisco is evolving for improved catalytic efficiency and CO2 assimilation in plants. Proc Natl Acad Sci. 2024 Mar 12;121(11):e2321050121.

11. Schulz L, Zarzycki J, Steinchen W, Hochberg GKA, Erb TJ. Layered entrenchment maintains essentiality in the evolution of Form I Rubisco complexes. EMBO J. 2024 Nov 18;1–12.

12. de Pins B, Greenspoon L, Bar-On YM, Shamshoum M, Ben-Nissan R, Milshtein E, et al. A systematic exploration of bacterial form I rubisco maximal carboxylation rates. EMBO J. 2024 July 15;43(14):3072–83.

13. Young JN, Heureux AMC, Sharwood RE, Rickaby REM, Morel FMM, Whitney SM. Large variation in the Rubisco kinetics of diatoms reveals diversity among their carbon-concentrating mechanisms. J Exp Bot. 2016 May 1;67(11):3445–56.

14. Lin MT, Occhialini A, Andralojc PJ, Parry MAJ, Hanson MR. A faster Rubisco with potential to increase photosynthesis in crops. Nature. 2014 Sept 1;513(7519):547–50.

15. Whitney SM, Baldet P, Hudson GS, Andrews TJ. Form I Rubiscos from non-green algae are expressed abundantly but not assembled in tobacco chloroplasts. Plant J. 2001;26(5):535–47.

16. Iñiguez C, Aguiló-Nicolau P, Galmés J. Improving photosynthesis through the enhancement of Rubisco carboxylation capacity. Biochem Soc Trans. 2021 Oct 8;49(5):2007–19.

17. Zhou Y, Whitney S. Directed Evolution of an Improved Rubisco; In Vitro Analyses to Decipher Fact from Fiction. Int J Mol Sci. 2019 Jan;20(20):5019.

18. Cai Z, Liu G, Zhang J, Li Y. Development of an activity-directed selection system enabled significant improvement of the carboxylation efficiency of Rubisco. Protein Cell. 2014 July 1;5(7):552–62.

19. Smith SA, Tabita FR. Positive and Negative Selection of Mutant Forms of Prokaryotic (Cyanobacterial) Ribulose-1,5-bisphosphate Carboxylase/Oxygenase. J Mol Biol. 2003 Aug 15;331(3):557–69.

20. Zhao L, Cai Z, Li Y, Zhang Y. Engineering Rubisco to enhance CO2 utilization. Synth Syst Biotechnol. 2024 Mar 1;9(1):55–68.

21. Davidi D, Shamshoum M, Guo Z, Bar-On YM, Prywes N, Oz A, et al. Highly active rubiscos discovered by systematic interrogation of natural sequence diversity. EMBO J. 2020 Sept 15;39(18):e104081.

22. McDonald JL, Shapiro NP, Mengiste AA, Kaines S, Whitney SM, Wilson RH, et al. In vivo directed evolution of an ultrafast Rubisco from a semianaerobic environment imparts oxygen resistance. Proc Natl Acad Sci. 2025 July 8;122(27):e2505083122.

23. Badger MR, Bek EJ. Multiple Rubisco forms in proteobacteria: their functional significance in relation to CO2 acquisition by the CBB cycle. J Exp Bot. 2008 May 1;59(7):1525–41.

24. Liu AK, Pereira JH, Kehl AJ, Rosenberg DJ, Orr DJ, Chu SKS, et al. Structural plasticity enables evolution and innovation of RuBisCO assemblies. Sci Adv. 2022 Aug 26;8(34):eadc9440.

25. Durão P, Aigner H, Nagy P, Mueller-Cajar O, Hartl FU, Hayer-Hartl M. Opposing effects of folding and assembly chaperones on evolvability of Rubisco. Nat Chem Biol. 2015 Feb;11(2):148–55.

26. Mao Y, Catherall E, Díaz-Ramos A, Greiff GRL, Azinas S, Gunn L, et al. The small subunit of Rubisco and its potential as an engineering target. J Exp Bot. 2023 Jan 11;74(2):543–61.

27. Lin MT, Salihovic H, Clark FK, Hanson MR. Improving the efficiency of Rubisco by resurrecting its ancestors in the family Solanaceae. Sci Adv. 2022 Apr 15;8(15):eabm6871.

28. Gunn LH, Martin Avila E, Birch R, Whitney SM. The dependency of red Rubisco on its cognate activase for enhancing plant photosynthesis and growth. Proc Natl Acad Sci. 2020 Oct 13;117(41):25890–6.

29. Prywes N, Phillips NR, Oltrogge LM, Lindner S, Taylor-Kearney LJ, Tsai YCC, et al. A map of the rubisco biochemical landscape. Nature [Internet]. 2025 Jan 22; Available from: 10.1038/s41586-024-08455-0

30. Prywes N, Phillips NR, Tuck OT, Valentin-Alvarado LE, Savage DF. Rubisco Function, Evolution, and Engineering. Annu Rev Biochem. 2023 June 20;92(1):385–410.

31. Satagopan S, Chan S, Perry LJ, Tabita FR. Structure-Function Studies with the Unique Hexameric Form II Ribulose-1,5-bisphosphate Carboxylase/Oxygenase (Rubisco) from Rhodopseudomonas palustris*. J Biol Chem. 2014 Aug 1;289(31):21433–50.

32. Zhang J, Zhao L, Liu G, Zhang Y, Cai Z, Li Y. A single amino acid substitution increases both carboxylation turnover number and CO2 affinity of form II Rubisco. Biochem Biophys Res Commun. 2025 July 1;768:151940.

33. Wilson RH, Martin-Avila E, Conlan C, Whitney SM. An improved Escherichia coli screen for Rubisco identifies a protein–protein interface that can enhance CO2-fixation kinetics. J Biol Chem. 2018 Jan 5;293(1):18–27.

34. Hong S, Spreitzer RJ. Complementing Substitutions at the Bottom of the Barrel Influence Catalysis and Stability of Ribulose-bisphosphate Carboxylase/Oxygenase*. J Biol Chem. 1997 Apr 25;272(17):11114–7.

35. Satagopan S, Scott SS, Smith TG, Tabita FR. A Rubisco Mutant That Confers Growth under a Normally “Inhibitory” Oxygen Concentration. Biochemistry. 2009 Sept 29;48(38):9076–83.

36. Satagopan S, Huening KA, Robert TF. Selection of Cyanobacterial (Synechococcus sp. Strain PCC 6301) RubisCO Variants with Improved Functional Properties That Confer Enhanced CO2-Dependent Growth of Rhodobacter capsulatus, a Photosynthetic Bacterium. mBio. 2019 July 23;10(4):10.1128/mbio.01537-19.

37. Wittmann BJ, Yue Y, Arnold FH. Informed training set design enables efficient machine learning-assisted directed protein evolution. Cell Syst. 2021 Nov 17;12(11):1026–1045.e7.

38. Chu HY, Fong JHC, Thean DGL, Zhou P, Fung FKC, Huang Y, et al. Accurate top protein variant discovery via low-N pick-and-validate machine learning. Cell Syst. 2024 Feb 21;15(2):193–203.e6.

39. Fowler DM, Fields S. Deep mutational scanning: a new style of protein science. Nat Methods. 2014 Aug;11(8):801–7.

40. Gantz M, Neun S, Medcalf EJ, van Vliet LD, Hollfelder F. Ultrahigh-Throughput Enzyme Engineering and Discovery in In Vitro Compartments. Chem Rev. 2023 May 10;123(9):5571–611.

41. Hoffmann UA, Schuppe AZ, Knave A, Sporre E, Brismar H, Englund E, et al. A Cyanobacterial Screening Platform for Rubisco Mutant Variants. ACS Synth Biol. 2025 July 18;14(7):2619–33.

42. Hagemann M, Hess WR. Systems and synthetic biology for the biotechnological application of cyanobacteria. Curr Opin Biotechnol. 2018 Feb 1;49:94–9.

43. Kim DS, Moreno-Cabezuelo JÁ, Schulz EN, Lea-Smith DJ, Sagaram US. Recent advances in engineering fast-growing cyanobacterial species for enhanced CO2 fixation. Front Clim [Internet]. 2024;6. Available from: https://www.frontiersin.org/journals/climate/articles/10.3389/fclim.2024.1412232

44. Hopf TA, Green AG, Schubert B, Mersmann S, Schärfe CPI, Ingraham JB, et al. The EVcouplings Python framework for coevolutionary sequence analysis. Bioinformatics. 2019 May 1;35(9):1582–4.

45. Hopf TA, Ingraham JB, Poelwijk FJ, Schärfe CPI, Springer M, Sander C, et al. Mutation effects predicted from sequence co-variation. Nat Biotechnol. 2017 Feb 1;35(2):128–35.

46. Biswas S, Khimulya G, Alley EC, Esvelt KM, Church GM. Low-N protein engineering with data-efficient deep learning. Nat Methods. 2021 Apr 1;18(4):389–96.

47. Riesselman AJ, Ingraham JB, Marks DS. Deep generative models of genetic variation capture the effects of mutations. Nat Methods. 2018 Oct 1;15(10):816–22.

48. Hsu C, Nisonoff H, Fannjiang C, Listgarten J. Learning protein fitness models from evolutionary and assay-labeled data. Nat Biotechnol. 2022 July 1;40(7):1114–22.

49. Notin P, Kollasch A, Ritter D, van Niekerk L, Paul S, Spinner H, et al. ProteinGym: Large-Scale Benchmarks for Protein Fitness Prediction and Design. In: Oh A, Naumann T, Globerson A, Saenko K, Hardt M, Levine S, editors. Advances in Neural Information Processing Systems [Internet]. Curran Associates, Inc.; 2023. p. 64331–79. Available from: https://proceedings.neurips.cc/paper_files/paper/2023/file/cac723e5ff29f65e3fcbb0739ae91bee-Paper-Datasets_and_Benchmarks.pdf

50. Bloom JD, Labthavikul ST, Otey CR, Arnold FH. Protein stability promotes evolvability. Proc Natl Acad Sci. 2006 Apr 11;103(15):5869–74.

51. Rizo J, Encarnación-Guevara S. Bacterial protein acetylation: mechanisms, functions, and methods for study. Front Cell Infect Microbiol [Internet]. 2024;14. Available from: https://www.frontiersin.org/journals/cellular-and-infection-microbiology/articles/10.3389/fcimb.2024.1408947

52. Wilson RH, Alonso H, Whitney SM. Evolving Methanococcoides burtonii archaeal Rubisco for improved photosynthesis and plant growth. Sci Rep. 2016 Mar 1;6(1):22284.

53. Wilson RH, Martin-Avila E, Conlan C, Whitney SM. An improved Escherichia coli screen for Rubisco identifies a protein–protein interface that can enhance CO2-fixation kinetics. J Biol Chem. 2018 Jan 5;293(1):18–27.

54. Jahn M, Vialas V, Karlsen J, Maddalo G, Edfors F, Forsström B, et al. Growth of Cyanobacteria Is Constrained by the Abundance of Light and Carbon Assimilation Proteins. Cell Rep. 2018 Oct 9;25(2):478–486.e8.

55. Amaral J, Lobo AKM, Carmo-Silva E. Regulation of Rubisco activity in crops. New Phytol. 2024;241(1):35–51.

56. Wittemeier L, Arrivault S, Neumann N, Maček B, Schmidt N, Hagemann M, et al. In vivo RUBISCO activity in Synechocystis is regulated by RuBP availability [Internet]. bioRxiv; 2025 [cited 2025 Nov 13]. p. 2025.06.06.658252. Available from: https://www.biorxiv.org/content/10.1101/2025.06.06.658252v1

57. Taylor TC, Andersson I. Structural transitions during activation and ligand binding in hexadecameric Rubisco inferred from the crystal structure of the activated unliganded spinach enzyme. Nat Struct Biol. 1996 Jan 1;3(1):95–101.

58. Duff AP, Andrews TJ, Curmi PMG. The transition between the open and closed states of rubisco is triggered by the inter-phosphate distance of the bound bisphosphate11Edited by D. Rees. J Mol Biol. 2000 May 19;298(5):903–16.

59. Satagopan S, North JA, Arbing MA, Varaljay VA, Haines SN, Wildenthal JA, et al. Structural Perturbations of Rhodopseudomonas palustris Form II RuBisCO Mutant Enzymes That Affect CO2 Fixation. Biochemistry. 2019 Sept 17;58(37):3880–92.

60. Li Z, Lanster DL, Badran AH. Synthetic approaches to enhance biological carbon capture. Curr Opin Biotechnol. 2025 Oct 1;95:103350.

61. Rao RM, Liu J, Verkuil R, Meier J, Canny J, Abbeel P, et al. MSA transformer. In PMLR; 2021. p. 8844–56.

62. Notin P, Weitzman R, Marks DS, Gal Y. ProteinNPT: Improving Protein Property Prediction and Design with Non-Parametric Transformers [Internet]. bioRxiv; 2023 [cited 2025 Nov 17]. p. 2023.12.06.570473. Available from: https://www.biorxiv.org/content/10.1101/2023.12.06.570473v1

63. Rao R, Liu J, Verkuil R, Meier J, Canny JF, Abbeel P, et al. MSA Transformer [Internet]. bioRxiv; 2021 [cited 2025 Nov 17]. p. 2021.02.12.430858. Available from: https://www.biorxiv.org/content/10.1101/2021.02.12.430858v3

64. Capó-Bauçà S, Font-Carrascosa M, Ribas-Carbó M, Pavlovič A, Galmés J. Biochemical and mesophyll diffusional limits to photosynthesis are determined by prey and root nutrient uptake in the carnivorous pitcher plant Nepenthes × ventrata. Ann Bot. 2020 June 19;126(1):25–37.

65. Kane H, Viil J, Entsch B, Paul K, Morell M, Andrews T. An Improved Method for Measuring the CO2/O2 Specificity of Ribulosebisphosphate Carboxylase-Oxygenase. Aust J Plant Physiol. 1994 Aug 1;21(4):449–61.

66. Kubien DS, Brown CM, Kane HJ. Quantifying the Amount and Activity of Rubisco in Leaves. In: Carpentier R, editor. Photosynthesis Research Protocols [Internet]. Totowa, NJ: Humana Press; 2011 [cited 2025 Nov 13]. p. 349–62. Available from: 10.1007/978-1-60761-925-3_27

67. McDonald JL, Shapiro NP, Whitney SM, Wilson RH, Shoulders MD. In vivo directed evolution of an ultra-fast RuBisCO from a semi-anaerobic environment imparts oxygen resistance [Internet]. bioRxiv; 2025 [cited 2025 Apr 25]. p. 2025.02.17.638297. Available from: https://www.biorxiv.org/content/10.1101/2025.02.17.638297v2

68. Lanster DL, Li Z, Badran AH. A Genetically Encoded System to Quantify and Evolve RuBisCO-Catalyzed Carbon Fixation [Internet]. bioRxiv; 2024 [cited 2025 Nov 13]. p. 2024.05.15.594374. Available from: https://www.biorxiv.org/content/10.1101/2024.05.15.594374v1

69. Cheng P, Mao C, Tang J, Yang S, Cheng Y, Wang W, et al. Zero-shot prediction of mutation effects with multimodal deep representation learning guides protein engineering. Cell Res. 2024 Sept;34(9):630–47.

70. van der Flier F, Estell D, Pricelius S, Dankmeyer L, van Stigt Thans S, Mulder H, et al. Enzyme structure correlates with variant effect predictability. Comput Struct Biotechnol J. 2024 Dec 1;23:3489–97.

71. Wu Z, Kan SBJ, Lewis RD, Wittmann BJ, Arnold FH. Machine learning-assisted directed protein evolution with combinatorial libraries. Proc Natl Acad Sci. 2019 Apr 30;116(18):8852–8.

72. Sun H, He L, Deng P, Liu G, Zhao Z, Jiang Y, et al. Accelerating protein engineering with fitness landscape modelling and reinforcement learning. Nat Mach Intell. 2025 Sept;7(9):1446–60.

73. Zhao L, Cai Z, Li Y, Zhang Y. Engineering Rubisco to enhance CO2 utilization. Synth Syst Biotechnol. 2024 Mar 1;9(1):55–68.

74. Zhou Y, Gunn LH, Birch R, Andersson I, Whitney SM. Grafting Rhodobacter sphaeroides with red algae Rubisco to accelerate catalysis and plant growth. Nat Plants. 2023 June;9(6):978–86.

75. Manning T, Birch R, Stevenson T, Nugent G, Whitney S. Bacterial Form II Rubisco can support wild-type growth and productivity in Solanum tuberosum cv. Desiree (potato) under elevated CO2. PNAS Nexus. 2023 Feb 1;2(2):pgac305.

76. Li H. Minimap2: pairwise alignment for nucleotide sequences. Bioinformatics. 2018 Sept 15;34(18):3094–100.

77. Li H, Handsaker B, Wysoker A, Fennell T, Ruan J, Homer N, et al. The Sequence Alignment/Map format and SAMtools. Bioinformatics. 2009 Aug 15;25(16):2078–9.

78. Martin M. Cutadapt removes adapter sequences from high-throughput sequencing reads. EMBnetjournal Vol 17 No 1 Gener Seq Data Anal [Internet]. 2011; Available from: https://journal.embnet.org/index.php/embnetjournal/article/view/200

79. Du W, Jongbloets JA, Pineda Hernández H, Bruggeman FJ, Hellingwerf KJ, Branco dos Santos F. Photonfluxostat: A method for light-limited batch cultivation of cyanobacteria at different, yet constant, growth rates. Algal Res. 2016 Dec 1;20:118–25.

80. Miao R, Jahn M, Shabestary K, Peltier G, Hudson EP. CRISPR interference screens reveal growth–robustness tradeoffs in *Synechocystis* sp. PCC 6803 across growth conditions. Plant Cell. 2023 July 26;koad208.

81. Langmead B, Salzberg SL. Fast gapped-read alignment with Bowtie 2. Nat Methods. 2012 Apr 1;9(4):357–9.

82. Love MI, Huber W, Anders S. Moderated estimation of fold change and dispersion for RNA-seq data with DESeq2. Genome Biol. 2014 Dec 5;15(12):550.

83. Miton CM, Tokuriki N. How mutational epistasis impairs predictability in protein evolution and design. Protein Sci. 2016 July 1;25(7):1260–72.

84. Madeira F, Madhusoodanan N, Lee J, Eusebi A, Niewielska A, Tivey ARN, et al. The EMBL-EBI Job Dispatcher sequence analysis tools framework in 2024. Nucleic Acids Res. 2024 July;52(W1):W521–5.

85. Sievers F, Higgins DG. Clustal Omega. Curr Protoc Bioinforma. 2014 Dec 1;48(1):3.13.1- 3.13.16.

86. Waterhouse AM, Procter JB, Martin DMA, Clamp M, Barton GJ. Jalview Version 2—a multiple sequence alignment editor and analysis workbench. Bioinformatics. 2009 May 1;25(9):1189–91.

87. Meng EC, Goddard TD, Pettersen EF, Couch GS, Pearson ZJ, Morris JH, et al. UCSF ChimeraX: Tools for structure building and analysis. Protein Sci. 2023 Nov 1;32(11):e4792.

88. Schrödinger, LLC. The PyMOL Molecular Graphics System, Version 1.8. 2015.

89. Tien MZ, Meyer AG, Sydykova DK, Spielman SJ, Wilke CO. Maximum Allowed Solvent Accessibilites of Residues in Proteins. PLOS ONE. 2013 Nov 21;8(11):e80635.

90. Abramson J, Adler J, Dunger J, Evans R, Green T, Pritzel A, et al. Accurate structure prediction of biomolecular interactions with AlphaFold 3. Nature. 2024 June;630(8016):493–500.

91. Marques SM, Borko S, Vavra O, Dvorsky J, Kohout P, Kabourek P, et al. Caver Web 2.0: analysis of tunnels and ligand transport in dynamic ensembles of proteins. Nucleic Acids Res. 2025 July 7;53(W1):W132–42.

